# Protein aggregation is a consequence of the dormancy-inducing membrane toxin TisB in *Escherichia coli*

**DOI:** 10.1101/2024.02.22.581605

**Authors:** Florian H. Leinberger, Liam Cassidy, Daniel Edelmann, Nicole E. Schmid, Patrick Blumenkamp, Sebastian Schmidt, Ana Natriashvili, Maximilian H. Ulbrich, Andreas Tholey, Hans-Georg Koch, Bork A. Berghoff

## Abstract

Bacterial dormancy is a valuable strategy to survive stressful conditions. Toxins from chromosomal toxin-antitoxin systems have the potential to halt cell growth, induce dormancy and eventually promote a stress-tolerant persister state. Due to their potential toxicity when overexpressed, sophisticated expression systems are needed when studying toxin genes. Here, we present an optimized plasmid expression system for toxin genes based on an artificial 5’ untranslated region. We applied the system to induce expression of the toxin gene *tisB* from the chromosomal type I toxin-antitoxin system *tisB/istR-1* in *Escherichia coli*. TisB is a small hydrophobic protein that targets the inner membrane, resulting in depolarization and ATP depletion. We analyzed TisB-producing cells by RNA-sequencing and revealed several genes with a role in recovery from TisB-induced dormancy, including the chaperone genes *ibpB*, *spy* and *cpxP*. The importance of chaperone genes suggested that TisB- producing cells are prone to protein aggregation, which was validated by an *in vivo* fluorescent reporter system. We moved on to show that TisB is an essential factor for protein aggregation upon DNA damage mediated by the fluoroquinolone antibiotic ciprofloxacin in *E. coli* wild-type cells. The occurrence of protein aggregates correlates with an extended dormancy duration, which underscores their importance for the life cycle of TisB-dependent persister cells.

**Importance:** Protein aggregates occur in all living cells due to misfolding of proteins. In bacteria, protein aggregation is associated with cellular inactivity, which is related to dormancy and tolerance to stressful conditions, including the exposure to antibiotics. In *Escherichia coli*, the membrane toxin TisB is an important factor for dormancy and antibiotic tolerance upon DNA damage mediated by the fluoroquinolone antibiotic ciprofloxacin. Here, we show that TisB provokes protein aggregation, which in turn promotes a deeper state of cellular dormancy. Our study suggests that protein aggregation is a consequence of membrane toxins with the potential to affect the duration of dormancy and the outcome of antibiotic therapy.

## Introduction

Bacteria constantly encounter stressful conditions due to sudden changes in their environments. They can tolerate these stress conditions to some extent and are able to maintain cellular integrity by switching on designated response systems, which sense these conditions and adjust the expression of specific genes to counteract the harmful situation. Under extreme hostile conditions, however, regular stress response systems may fail to protect cells from lethal damages, a situation that is unpredictable and demands alternative survival strategies. One possibility is the formation of dormant cells, which are characterized by reduced cellular activity, growth arrest, and the ability to withstand even extreme stress conditions (1, 2). Dormancy typically occurs only in a fraction of cells and, therefore, represents an example of phenotypic heterogeneity that is considered as a bet-hedging strategy for survival in unpredictable environments: some bacteria sacrifice their own propagation to ensure continuity of the genotype in case of extreme hostile conditions (3).

Bacterial dormancy occurs in different shapes and degrees. On the one hand, bacteria may establish extreme morphotypes, such as myxospores within fruiting bodies of myxobacteria or endospores of some gram-positive bacteria, which may reside in a dormant state for many years (4, 5). On the other hand, bacterial populations almost constantly generate cells that are morphologically similar to their siblings, but have entered a transient state of reduced activity from which they can rapidly recover. A prominent example are so-called persister cells, which are well known for their ability to survive antibiotic treatments (6–8). They have gained increasing attention as they may cause infection relapse or serve as a reservoir for antibiotic resistance development (9–11). As it stands right now, there are many ways into the persister state, including spontaneous events (12, 13), nutrient limitation and starvation (14, 15), metabolic perturbations (16), oxidative stress (17, 18), and low energy levels (19, 20). However, it is not entirely clear whether a combination of these events is necessary to reduce cellular activity to such an extent that persister formation is promoted. In this respect, not every dormant cell is a persister cell, but dormancy increases the likelihood to reach the persister state (21).

Another possibility to induce dormancy are toxin-antitoxin (TA) systems, which are ubiquitously found in bacteria and contribute to stress responses or stabilization of mobile genetic elements (22, 23). Different TA system types have been identified, but they all have in common that the antitoxin inhibits toxin activity or prevents toxin production (22–24), which likely restricts toxin-dependent effects to specific (stress) conditions (25, 26). Whether or not toxins from TA systems induce a persister state is subject to current debate (27, 28), but the contribution of toxins to bacterial dormancy and condition- dependent persister formation seems plausible (26, 29–31). One well-studied toxin with a potential influence on dormancy and persistence is TisB from the type I TA system *tisB/istR-1* in *Escherichia coli* (32–34). TisB is a small hydrophobic protein that targets the inner membrane and leads to membrane depolarization, ATP depletion, and further secondary effects, such as reactive oxygen species formation and inhibition of translation (25, 29, 35–37). The reduced energy level in TisB-producing cells is expected to support persister formation, especially under conditions of DNA damage, when the corresponding *tisB* toxin gene is strongly induced upon auto-cleavage of the LexA repressor as part of the SOS response (25, 31, 38, 39). However, transcription of *tisB* is not sufficient to produce the TisB protein, because the primary *tisB* mRNA (+1 mRNA) is translationally inactive due to an inhibitory secondary structure in its 5’ part. The +1 mRNA needs to undergo processing into the active +42 mRNA to be translated (40). Translation of the +42 mRNA depends on a non-canonical translation initiation mechanism that involves a single-stranded ribosome standby site (RSS), a 5’ pseudoknot structure, and ribosomal protein S1 (41, 42). However, translation of +42 mRNA is efficiently inhibited by the RNA antitoxin IstR-1 (39, 40). Hence, two regulatory RNA elements (secondary structure in the +1 mRNA and antitoxin IstR-1) limit *tisB* expression to SOS conditions and set a threshold for TisB production in individual cells, thereby causing phenotypic heterogeneity (25, 43).

An early transcriptome study demonstrated that heterologous production of TisB and other membrane toxins led to the induction of several stress response genes (44), indicating that these toxins cause stress due to primary and secondary effects (29, 37). However, heterologous toxin expression systems tend to produce excessive effects. In the current study, we aimed at constructing an optimized expression system to study the TisB-dependent stress response. Here, we observed that moderate *tisB* expression elicits a stress response that contributes to recovery from TisB-induced dormancy. Up- regulation of several chaperone genes suggested that TisB provokes protein aggregation, which was validated by using a fluorescent reporter system. Intriguingly, we found that the DNA-damaging antibiotic ciprofloxacin causes protein aggregation in a TisB-dependent manner, and that protein aggregates affect the dormancy depth of persister cells. Our study supports the view that TisB – and probably other type I toxins – affect dormancy and persistence through a variety of downstream effects, including protein aggregation.

## Results

### An improved expression system for investigation of TisB-induced dormancy

Production of the membrane toxin TisB from the type I TA system *tisB/istR-1* inflicts a stressful situation, including perturbation of membrane functioning, energy depletion and further secondary effects (29, 45). Recent work on TisB has highlighted the importance of particular stress-related proteins in the context of TisB-dependent persistence, such as superoxide dismutases and alkyl hydroperoxide reductase (37, 46). In order to grasp the global response to TisB-mediated stress, we aimed at constructing an inducible expression system that provokes TisB-dependent effects but avoids high TisB levels and concomitant TisB toxicity (35, 47). In *E. coli*, pBAD plasmids are applied for controllable gene expression from the P_BAD_ promoter using L-arabinose (L-ara) as inducer. When using the pBAD derivative p+42-*tisB* (35), transcription from the P_BAD_ promoter produces the native *tisB* +42 mRNA, which is translationally active due to its accessible RSS and the existence of a Shine-Dalgarno (SD) sequence (Fig. 1A). However, *tisB* induction from p+42-*tisB* reduces the number of colony forming units (CFU) by at least 10-fold, indicating TisB toxicity and probably cell death (35). Since *tisB* expression from its chromosomal gene copy is not expected to cause cell death (35), but rather supports stabilization of a growth-arrested state (29), p+42-*tisB* does not represent a suitable expression system to study authentic TisB effects. Expression strength can be modulated by plasmid copy number and promoter strength (48). Alternatively, the efficiency of translation can be modulated. We followed the latter strategy and replaced the native *tisB* upstream region by a short artificial region with a length of 20 bp, lacking a SD sequence (49) (Fig. 1A). The resulting plasmid, termed p0SD-*tisB*, was transferred to *E. coli* wild type MG1655. Using 3xFLAG fusions and Western blot analysis, we compared the p+42- *tisB* and p0SD-*tisB* systems by assessing 3xFLAG-TisB protein levels (Fig. 1B). 3xFLAG-TisB levels were reduced by ∼10-fold using the p0SD-*tisB* system, which was presumably due to a lower efficiency of translation initiation, but might also be partly attributable to lower steady-state levels of the *0SD- 3xFLAG-tisB* mRNA (Fig. S1). Optical density (OD_600_) measurements demonstrated that TisB induction from plasmid p0SD-*tisB* by L-ara was sufficient to halt cell growth during exponential phase, while an empty pBAD control showed normal growth (Fig. 1C). There was, however, a short delay for growth inhibition with p0SD-*tisB* when compared to p+42-*tisB*. The primary effect of TisB is depolarization of the inner membrane (25, 36). We assessed depolarization by the potential-sensitive probe DiBAC_4_(3) after one hour of L-ara treatment during exponential phase. As expected, TisB expression from both p+42-*tisB* and p0SD-*tisB* caused an increase of intracellular DiBAC_4_(3) fluorescence compared to the empty pBAD control (Fig. 1D). However, depolarization was more pronounced with the p+42-*tisB* system, which was in agreement with higher TisB protein levels (Fig. 1B). Importantly, after one hour of L-ara treatment, 66% of cells were able to form colonies with the p0SD-*tisB* system, while this value dropped to 1% with p+42-*tisB* (Fig. 1E). These findings indicate that p0SD-*tisB* largely avoids TisB toxicity and, therefore, represents an optimal expression system to study TisB-induced dormancy.

**Figure 1.**
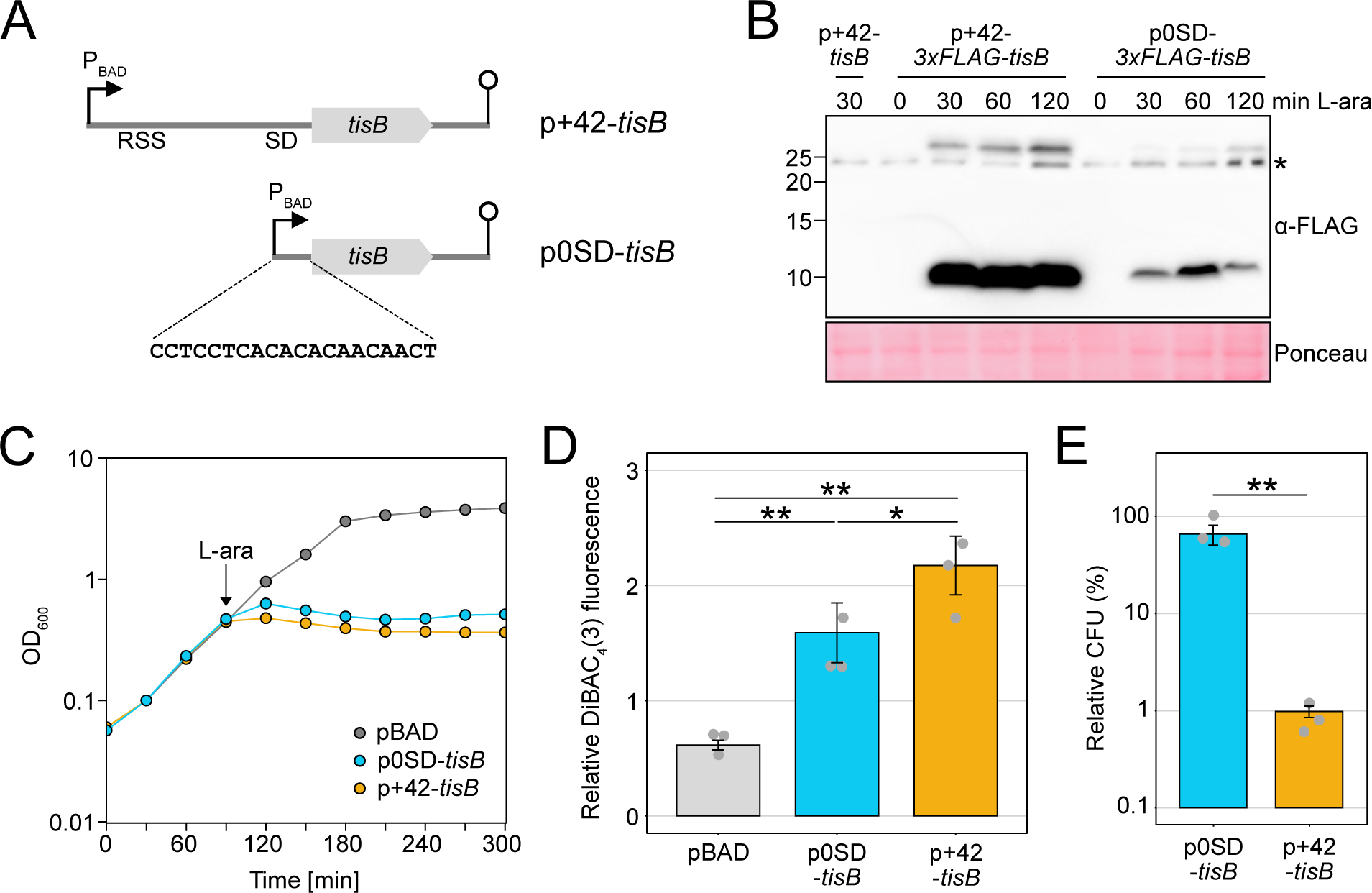
Characterization of an optimized *tisB* expression system. **(A)** Schematic representation of different *tisB* expression systems. The p+42-*tisB* plasmid contains the native *tisB* 5’ UTR, including a ribosome standby site (RSS) and a Shine-Dalgarno (SD) sequence. Transcription from the P_BAD_ promoter starts at the *tisB* +42 position. The p0SD-*tisB* plasmid contains the *tisB* coding sequence preceded by an artificial 20-bp 5’ UTR. Lollipop structures indicate Rho- independent terminators. **(B)** Detection of 3xFLAG-TisB. Wild type MG1655 harboring 3xFLAG-tag variants of p+42-*tisB* and p0SD-*tisB* were grown to an OD_600_ of ∼0.4 (exponential phase) and treated with L-ara (0.2%). Samples were collected at the indicated time points. Total protein was separated using Tricine-SDS-PAGE and transferred to PVDF membranes by electro-blotting. 3xFLAG-TisB was detected using an HRP-conjugated monoclonal α-FLAG antibody. As negative control, p+42-*tisB* was used. Two TisB-specific bands are visible, one at ∼10 kDa and one above 25 kDa. The asterisk indicates an unspecific band. Ponceau staining is shown as loading control. **(C)** Growth inhibition by TisB. Wild type MG1655, harboring p0SD-*tisB*, p+42-*tisB* or an empty pBAD plasmid, was treated with the inducer L-ara (0.2%) at an OD_600_ of ∼0.4 (exponential phase; arrow). The OD_600_ was measured over time. Data points indicate the mean of three biological replicates. **(D)** TisB-dependent membrane depolarization. Wild type MG1655 cells, harboring p0SD-*tisB*, p+42-*tisB* or an empty pBAD plasmid, were treated with the inducer L-ara (0.2%) for 1 hour when an OD_600_ of ∼0.4 was reached (exponential phase). Staining with the potential-sensitive probe DiBAC_4_(3) was applied to assess depolarization. The relative DiBAC_4_(3) fluorescence indicates the ratio between 1-hour and 0-hour measurements. Bars represent the mean of three biological replicates and error bars indicate the standard deviation. Dots show individual data points. ANOVA with post-hoc Tukey HSD test was performed (* p<0.05, ** p<0.01). **(E)** TisB toxicity with different expression systems. Wild type MG1655, harboring p0SD-*tisB* or p+42-*tisB*, was treated with L-ara (0.2%) during exponential phase (OD_600_ ∼0.4) for 1 hour. Pre- and post- treatment samples were used to determine relative CFU (%) by plating and counting. Bars represent the mean of three biological replicates and error bars indicate the standard deviation. Dots show individual data points. ANOVA with post-hoc Tukey HSD test was performed (** p<0.01).

### Cell populations adapt to moderate *tisB* expression

Elevated toxin levels were shown to increase phenotypic heterogeneity with respect to growth-arrest duration and persistence time (25, 50). The duration of toxin-induced growth arrest is reflected by the time that is needed by single cells to form colonies on agar plates, which can be quantified using the ScanLag method (51, 52). When *E. coli* wild type MG1655, containing p0SD-*tisB*, was grown to an OD_600_ of ∼0.4 (exponential phase) and plated on regular LB agar plates without L-ara (T0; Fig. 2A), the median colony appearance time was 820 min (Fig. 2B). The narrow appearance-time distribution indicated homogeneous lag times, as expected from exponentially growing populations. By contrast, when cultures were treated with L-ara for 30 min to induce TisB-dependent growth arrest before cells were spread on agar plates (T30; Fig. 2A), the median colony appearance time shifted to 1,120 min. In other words, cells needed on average 5 hours longer to form colonies. Furthermore, heterogeneity of colony appearance was clearly increased (Fig. 2B). Since the speed of colony growth could not account for the 5-hours shift (Fig. S2), we concluded that TisB production from the p0SD-*tisB* system generated populations with very heterogeneous growth-arrest durations. While results for a 60-min L-ara treatment (T60) were comparable to the 30-min time point, the median colony appearance time was only 960 min after 120 min of L-ara treatment (T120), indicating that heterogeneity declined over time (Fig. 2B). Even more intriguingly, relative CFU counts stayed at ∼50% during the first 60 min of L-ara treatment but increased to more than 90% after 120 min (Fig. 2C). Hence, cells regained their ability to form colonies at later stages. Together with the shorter growth-arrest duration observed for the 120-min time point, these data suggested that cell populations adapted to TisB production during the first two hours of toxin-mediated stress.

**Figure 2.**
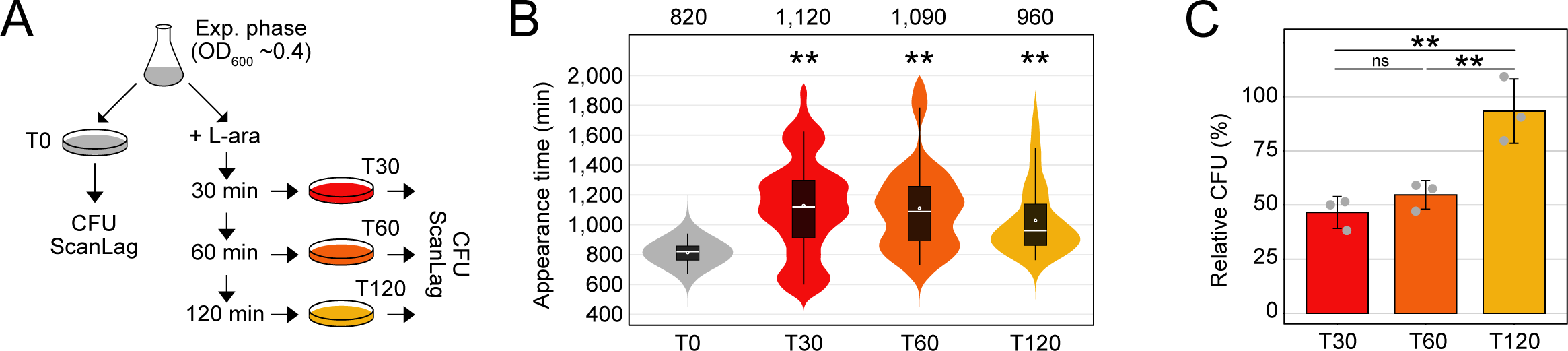
Cells adapt to moderate *tisB* expression. **(A)** Schematic representation of the performed experiment. Wild type MG1655, harboring the p0SD- *tisB* plasmid, was treated with L-ara (0.2%) in exponential phase (OD_600_ ∼0.4). At the indicated time points (T30, T60 and T120), cells were plated on LB agar without L-ara and colony growth was analyzed using the ScanLag method (see *Material and Methods*). As a control, cells were analyzed before L-ara was added (T0). **(B)** ScanLag analysis was applied to determine the colony appearance time after *tisB* expression. For each time point, colony appearance times are illustrated as violin box plots. Colonies from three biological replicates were combined (T0: n=154; T30: n=59; T60: n=103; T120: n=124). The white dot indicates the mean. The respective median appearance time (white bar) is shown on top of each plot. L-ara treated samples were compared to the control (T0) using a pairwise Wilcoxon rank sum test (** p<0.0001). **(C)** Colony counts increase upon progressing *tisB* expression. LB agar plates from (B) were used to determine colony counts. Pre- (T0) and post-treatment (T30, T60, T120) samples were used to determine relative CFU (%). Bars represent the mean of three biological replicates and error bars indicate the standard deviation. Dots show individual data points. ANOVA with post-hoc Tukey HSD test was performed (** p<0.01; ns: not significant).

### A global transcriptome analysis reveals TisB-dependent up-regulation of stress-related genes

We assumed that adaptation to TisB was driven by cellular stress responses. It has already been observed that type I toxins cause up-regulation of several stress-related genes (44), and we have shown that *tisB* expression provokes superoxide formation and up-regulation of *soxS* and the SoxRS regulon (37). To reveal the response to TisB on a global scale, transcriptome analysis of MG1655 p0SD- *tisB* was performed. Cultures were grown to an OD_600_ of ∼0.4 (exponential phase) and treated with L- ara for 30 min. Samples before and after L-ara treatment were collected and analyzed by RNA-seq, which identified 67 up-regulated and 66 down-regulated genes (log_2_ fold-change >2 and <‒2, *p*-value <0.01; Dataset S1). We specifically focused on up-regulated genes, as they might represent an active response to TisB. As expected, *tisB* and *soxS* were among the genes with the strongest up-regulation (Fig. 3A). To select candidates for further analysis, we compared the set of up-regulated genes from our RNA-seq analysis to (i) microarray data of heterologous *tisB* expression (44), (ii) proteome data of a de-regulated *tisB* strain (46), and (iii) transcriptional regulation data from the RegulonDB database (53). The regulon analysis highlighted genes that are transcriptionally regulated by CpxR, the response regulator from the CpxAR two-component system (Fig. S3). The Cpx system belongs to the envelope stress response and is mainly involved in sensing misfolded proteins in the inner membrane and periplasm (54). In total, we selected four CpxR-dependent genes: *cpxP*, *spy*, *yebE* and *yqaE* (Fig. 3A). Importantly, *cpxP*, *spy* and *yebE* were found in the transcriptome study by Fozo *et al.* (44), and *spy* and *yebE* were also found in the proteome study by Spanka *et al.* (46). CpxP and Spy are located in the periplasm and have chaperone functions; YebE and YqaE are poorly characterized inner membrane proteins. In addition, we selected *ydjM* (Fig. 3A), encoding another poorly characterized inner membrane protein. Like *tisB*, *ydjM* belongs to the LexA regulon and might have an important function during the SOS response. The transcriptome study by Fozo *et al.* showed that the *ibpAB* operon is upregulated upon type I toxin expression (44). Both *ibpA* and *ibpB* encode small heat-shock proteins (sHSPs) with chaperone functions in the cytoplasm. Since *ibpB* showed stronger induction than *ibpA* in our RNA-seq data (Dataset S1), we selected *ibpB* for further analysis (Fig. 3A). We applied quantitative reverse transcription PCR (qRT-PCR) to verify TisB-dependent induction of the selected genes. *soxS* and *tisB* were used as positive controls (Fig. 3B). To exclude that up-regulation of stress-related genes was due to the L-ara treatment, wild type MG1655 containing an empty pBAD plasmid was analyzed by qRT-PCR, clearly showing that L-ara alone was not sufficient to cause induction of the stress-related genes (Fig. 3B). Finally, *bhsA* and *yhcN* were selected, because they were among the genes with the strongest upregulation (Fig. 3A). Both genes encode DUF1471 domain-containing proteins that are located in the cell envelope and have a putative role in stress responses and/ or biofilm formation (55–57). Since qRT-PCR analysis did not produce reliable results for *bhsA* and *yhcN*, Northern blot analysis was performed, which confirmed their TisB-dependent induction (Fig. 3C and D). We note, however, that *tisB* expression caused accumulation of several mRNA degradation products, which was particularly evident for *yhcN* (Fig. 3D). Strong *tisB* expression causes rRNA degradation in less than one hour (35, 37, 47), but this was not observed when using the p0SD-*tisB* system (Fig. S1), suggesting that global RNA decay cannot account for *bhsA* and *yhcN* degradation. Degradation of *bhsA* and *yhcN* might have a biological function, such as generation of regulatory RNAs, but this needs further investigation.

**Figure 3.**
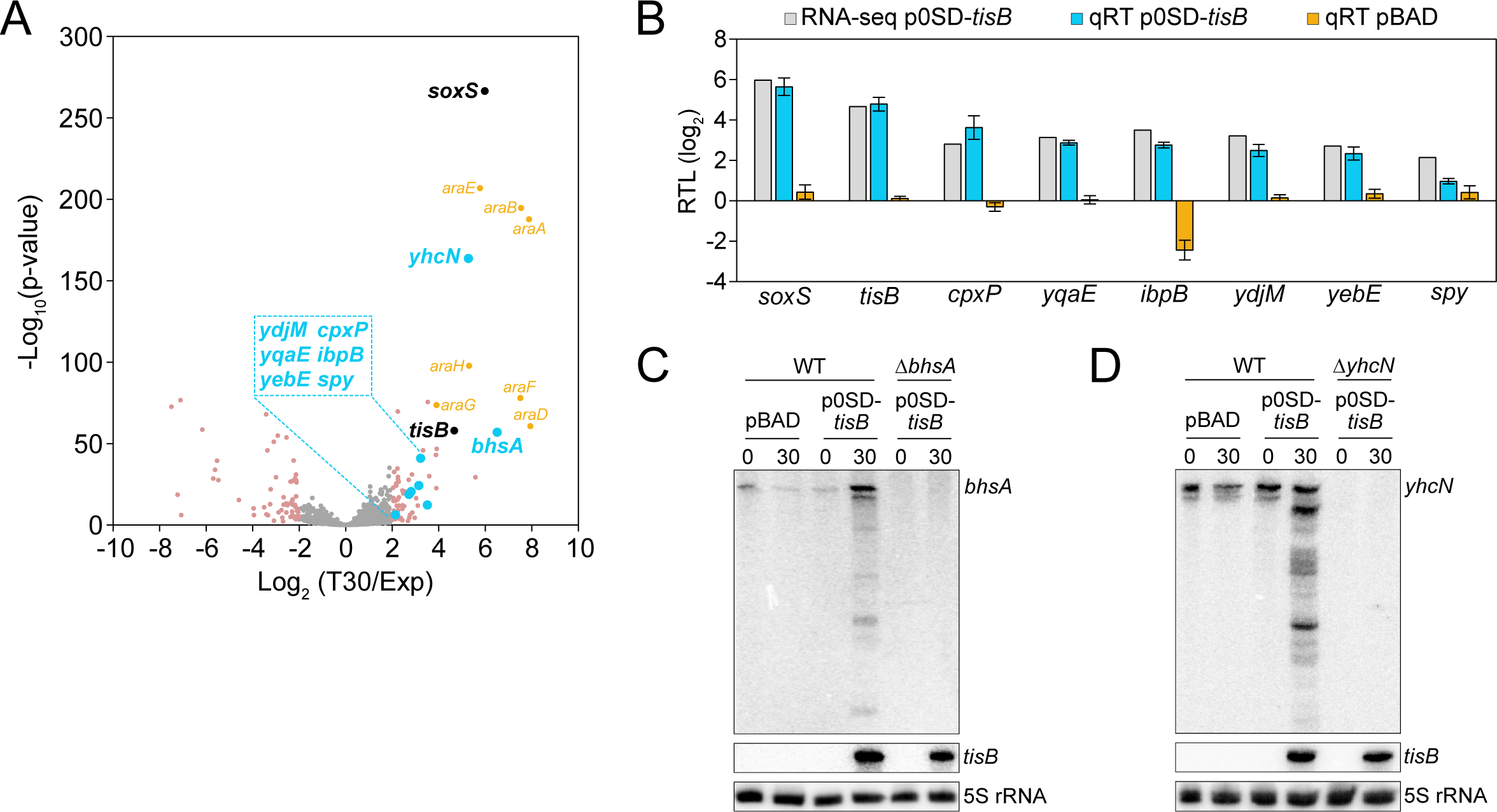
Identification of TisB-responsive genes by RNA-seq. **(A)** Global response to *tisB* expression. Wild type MG1655, harboring the p0SD-tisB plasmid, was treated with L-ara (0.2%) during exponential phase (OD_600_ ∼0.4) for 30 min. RNA samples extracted before (Exp) and after treatment (T30) were analyzed using RNA-seq. The volcano plot illustrates the log_2_ fold-change on the x-axis and the ‒log_10_(p-value) on the y-axis. Differentially expressed genes (log_2_ fold-change >2 or <‒2, p-value <0.01) are shown in pink. Selected candidates are highlighted in blue, while genes affected by L-ara are shown in orange (*araBAD*, *araE*, *araFGH*), and *tisB* and *soxS* are shown in black. **(B)** Confirmation of RNA-seq using qRT-PCR. Wild type MG1655, harboring p0SD-*tisB* (blue bars) or an empty pBAD plasmid (orange bars), was treated with L-ara (0.2%) during exponential phase (OD_600_ ∼0.4) for 30 min. Relative transcript levels (RTL; log_2_) were assessed by qRT-PCR (qRT). Log_2_ fold- changes from the RNA-seq analysis are shown for comparison (grey bars). Bars represent the mean of three biological replicates, with two technical replicates each, and error bars indicate the standard deviation. **(C, D)** Confirmation of RNA-seq using Northern blot analysis. Wild type MG1655, harboring p0SD-*tisB* or an empty pBAD plasmid, was treated with L-ara (0.2%) during exponential phase (OD_600_ ∼0.4) for 30 min. Total RNA was separated using urea-polyacrylamide gels and blotted onto nylon membranes. Radioactive probes binding to the coding region of **(C)** *bhsA* or **(D)** *yhcN* were applied for detection of transcripts. Corresponding deletion mutants (Δ*bhsA* or Δ*yhcN*) were used to show specificity of the probes. A *tisB* probe was applied to verify *tisB* induction from p0SD-*tisB*, and 5S rRNA was probed as loading control.

### Stress-related genes support recovery from TisB-induced dormancy

To evaluate the function of the selected candidates with respect to TisB-induced dormancy, we deleted the corresponding genes and transferred the p0SD-*tisB* plasmid to the resulting mutants. As expected, all mutants showed L-ara induced and TisB-dependent growth inhibition (Fig. S4). In a subsequent experiment, mutants were grown to an OD_600_ of ∼0.4 (exponential phase) and tested for their ability to form colonies after one hour of L-ara treatment. The relative CFU counts for the mutants ranged between 43% and 111%, which was not strikingly different when compared to the wild type (70%; Fig. 4A). We concluded that each individual gene only had a minor influence on the ability of TisB-producing cells to form colonies on LB agar plates. We reasoned that the stress-related genes might rather influence the growth-arrest duration by supporting the recovery from TisB-mediated stress (37, 46). Indeed, when using ScanLag, seven out of eight mutants showed a delayed recovery and significantly increased growth-arrest duration in comparison to the wild type, with Δ*bhsA* being the only exception (Fig. 4B). The growth-arrest duration, as measured by the colony appearance time, was prolonged by at least 80 min (Δ*ibpB*) and up to 220 min (Δ*yebE* and Δ*cpxP*). To exclude that the gene deletion itself and/ or the L-ara treatment would affect the colony appearance time, an empty pBAD plasmid was transferred to the mutants. The resulting strains were grown to exponential phase, treated with L-ara for one hour and analyzed by ScanLag. In this control experiment, none of the mutants showed a delayed colony appearance in comparison to the wild type (Fig. S5), clearly indicating that the stress-related genes have a particular function upon TisB-mediated stress and probably support the recovery process. Since four of the eight candidates belong to the CpxR regulon (Fig. 4A), we constructed a *cpxR* deletion mutant, transferred the p0SD-*tisB* plasmid, and induced *tisB* expression by L-ara. However, neither CFU counts nor colony appearance were significantly different in the *cpxR* mutant when compared to the wild type (Fig. S6), suggesting that induction of the CpxR regulon is not mandatory for recovery from TisB-mediated stress.

**Figure 4.**
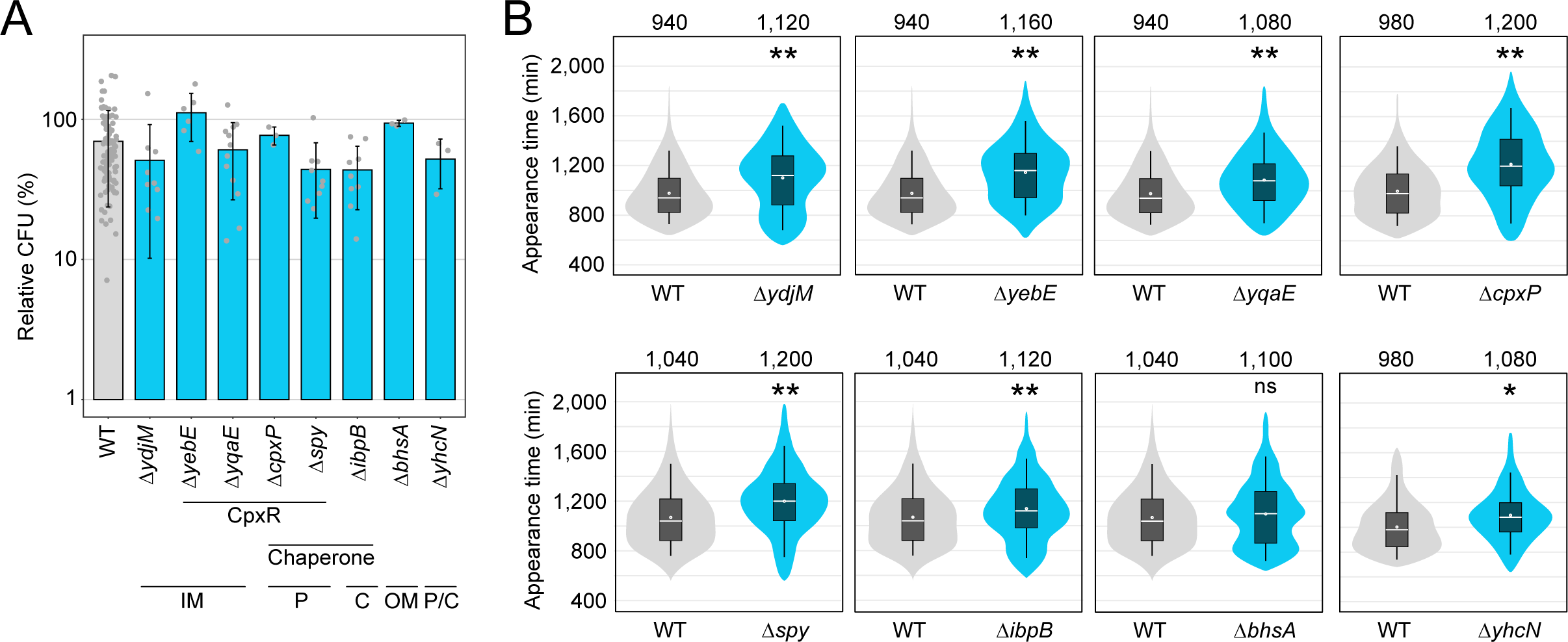
TisB-responsive genes mainly affect the recovery after *tisB* expression. **(A)** TisB toxicity in selected deletion mutants. Wild type (WT) MG1655 and deletion mutants, harboring the p0SD-*tisB* plasmid, were treated with L-ara (0.2%) during exponential phase (OD_600_ ∼0.4) for 1 hour. Pre- and post-treatment samples were used to determine relative CFU (%) by plating and counting. Bars represent the mean of at least three biological replicates and error bars indicate the standard deviation. Dots show individual data points (WT: n=102; Δ*ydjM*: n=9; Δ*yebE*: n=6; Δ*yqaE*: n=12; Δ*cpxP*: n=3; Δ*spy*: n=9; Δ*ibpB*: n=9; Δ*bhsA*: n=3; Δ*yhcN*: n=3). ANOVA with post-hoc Tukey HSD was performed (no significant difference between deletion mutants and the wild type was detected). It is indicated whether the genes are CpxR-dependent or have a chaperone activity. Their proposed cellular localization is given (C: cytoplasm, IM: inner membrane, P: periplasm, OM: outer membrane). **(B)** ScanLag analysis of selected deletion mutants. Wild type (WT) MG1655 and deletion mutants, harboring the p0SD-*tisB* plasmid, were treated with L-ara (0.2%) during exponential phase (OD_600_ ∼0.4) for 1 hour. ScanLag was applied to determine the colony appearance time after *tisB* expression. For each deletion mutant, colony appearance times are illustrated as violin box plots and compared to a corresponding wild type. Colonies from at least three biological replicates were combined (WT: n≥192; Δ*ydjM*: n=452; Δ*yebE*: n=383; Δ*yqaE*: n=393; Δ*cpxP*: n=252; Δ*spy*: n=356; Δ*ibpB*: n=682; Δ*bhsA*: n=365; Δ*yhcN*: n=192). The white dot indicates the mean. The respective median appearance time (white bar) is shown on top of each plot. Deletion mutants were compared to wild type MG1655 using a pairwise Wilcoxon rank sum test (* p<0.001, ** p<0.0001, ns: not significant). It should be noted that ScanLag results vary between individuals runs. For every mutant, statistical testing refers to the corresponding control strain (WT) from the same experimental run.

### TisB causes intracellular ATP depletion and protein aggregation

The importance of proteins with chaperone activity during recovery from TisB-induced growth arrest (Fig. 4) suggested that unfolded proteins and protein aggregates impose a challenge for TisB-producing cells. It was previously demonstrated that due to ATP depletion protein aggregates form and affect the dormancy of bacterial cells (58, 59). Since TisB is expected to decrease the intracellular ATP concentration due to depolarization of the inner membrane and breakdown of the proton motive force (31, 33, 35), intracellular ATP concentrations were measured before and after L-ara treatment in wild type MG1655 containing either p0SD-*tisB* or an empty pBAD plasmid. In the TisB-producing strain, a 60-min treatment with L-ara caused a ∼32-fold ATP reduction, while the ATP concentration remained unchanged in the control strain (Fig. 5A). To assess cytosolic protein aggregation as a likely consequence of ATP depletion, we applied a reporter strain that chromosomally expresses a monomeric superfolder green fluorescent protein (msGFP) fused to the C-terminus of the sHSP IbpA (60). As expected, cytosolic msGFP fluorescence changed from a diffuse to a punctuated pattern (i.e., formation of foci) after 15 min of heat shock at 47°C (Fig. S7). Since IbpA localizes to protein aggregates, the msGFP foci clearly indicated the formation of protein aggregates in the cytoplasm due to elevated temperature (59, 60). We performed a U-Net analysis (61) to count msGFP foci in individual cells (Fig. S7). Expression of *tisB* from p0SD-*tisB* in the *ibpA-msGFP* reporter background (60-min L-ara treatment) led to the formation of foci, with ∼48% of cells having three foci and ∼20% having two or four foci (Fig. 5B and C). As a control, the empty pBAD plasmid was transferred to the *ibpA-msGFP* reporter background, and the resulting strain was treated with L-ara. However, L-ara alone was not sufficient to cause foci formation (Fig. 5B and C). In order to demonstrate that functional TisB was needed for ATP depletion and foci formation, plasmid p0SD-*tisB-K12L* was applied for production of the TisB-K12L variant. TisB-K12L has the central lysine 12 replaced with leucine, leading to attenuated TisB activity without affecting membrane localization (37). As expected, TisB-K12L did not cause major ATP depletion (Fig. 5A). More intriguingly, a reporter strain containing p0SD-*tisB-K12L* displayed mainly cells without foci (∼83%) after 60 min of L-ara treatment (Fig. 5B and C). This control experiment demonstrated that production of a small membrane protein (TisB-K12L) is not sufficient to cause cytosolic protein aggregation, but rather that functional TisB toxin triggers the formation of protein aggregates, probably due to strong intracellular ATP depletion.

**Figure 5.**
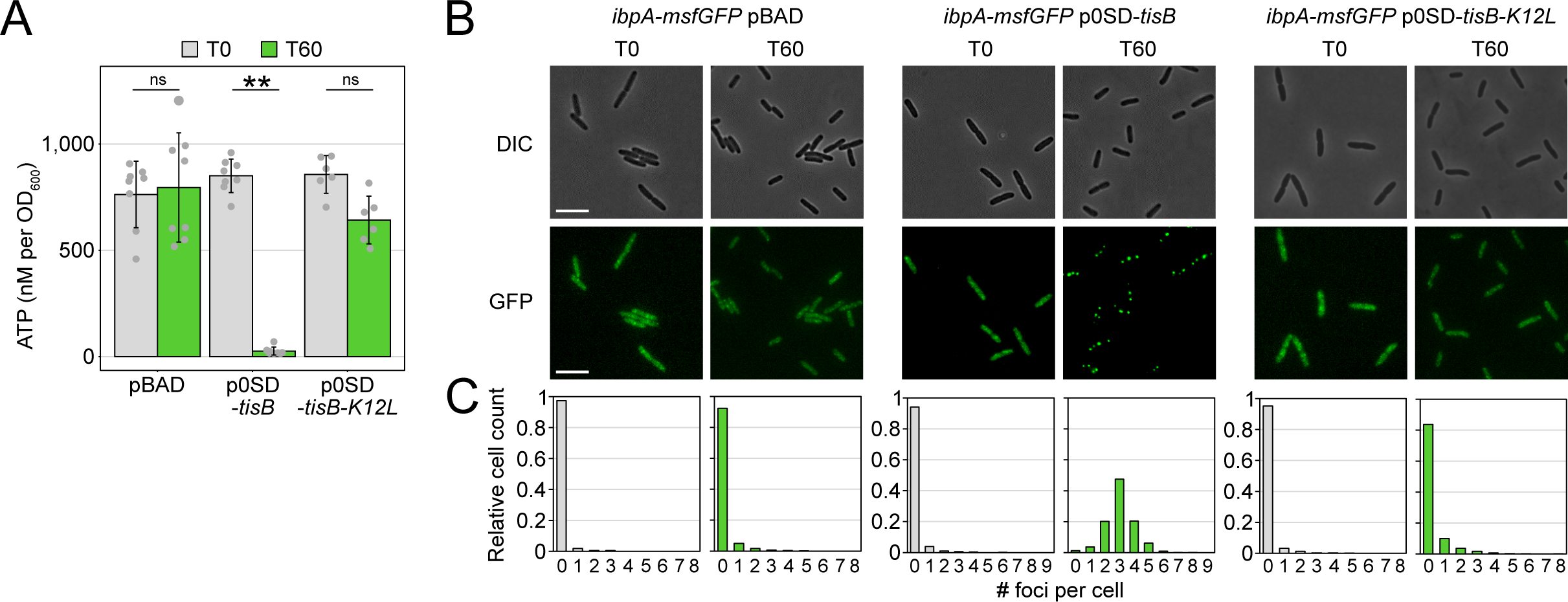
Expression of *tisB* causes cytoplasmic protein aggregation. **(A)** TisB-dependent ATP depletion. Wild type MG1655, harboring either an empty pBAD plasmid, the p0SD-*tisB* plasmid or the p0SD-*tisB-K12L* variant, was treated with L-ara (0.2%) during exponential phase (OD_600_ ∼0.4) for 60 min. A luciferase-based assay was applied to measure cellular ATP levels (nM per OD_600_) before (T0) and after L-ara treatment (T60). Bars represent the mean of at least six biological replicates and error bars indicate the standard deviation. Dots show individual data points (pBAD: n=8; p0SD-*tisB*: n=8; p0SD-*tisB-K12L*: n=6). ANOVA with post-hoc Tukey HSD test was performed (** p<0.01; ns: not significant). **(B)** TisB-dependent protein aggregation in the cytoplasm. Strain MG1655 *ibpA- msfGFP*, harboring either an empty pBAD plasmid, the p0SD-*tisB* plasmid or the p0SD-*tisB-K12L* variant, was treated with L-ara (0.2%) during exponential phase (T0; OD_600_ ∼0.4) for 60 min (T60). Differential interference contrast (DIC) images are displayed together with corresponding fluorescence images (GFP). White bars represent a length scale of 2 µm. Representative images from three biological replicates are shown. **(C)** Quantification of msGFP foci. The experiment shown in (B) was performed in three biological replicates. All images were evaluated using a U-Net neural network analysis and in-house image processing tools to automatically count msGFP foci per cell. At least 507 cells were analyzed for each condition (pBAD T0: n=507; pBAD T60: n=3019; p0SD-*tisB* T0: n=730; p0SD-*tisB* T60: n=1474; p0SD-*tisB-K12L* T0: n=1405; p0SD-*tisB-K12L* T60: n=1896).

### Ciprofloxacin provokes TisB-dependent protein aggregation

So far, we have shown that *tisB* expression from plasmid p0SD-*tisB* induces several stress-related genes, encoding – amongst others – the chaperones CpxP, Spy and IbpB. Deletion of these genes delays recovery of cells following TisB-mediated stress. Furthermore, we have observed strong ATP depletion and protein aggregation upon *tisB* expression from plasmid p0SD-*tisB*. While these experiments are helpful to appreciate the cellular consequences of *tisB* expression, they do not provide direct evidence for the consequences of *tisB* expression in wild-type cells. In wild-type cells, *tisB* transcription is strongly induced upon DNA damage through UV light or DNA-damaging agents, such as mitomycin C or ciprofloxacin (25, 38, 39, 62). When using the gyrase inhibitor ciprofloxacin (CIP), most TisB- dependent effects are observed only after approximately three hours of a high-dose treatment (29). We, therefore, treated wild type MG1655 and a corresponding *tisB* deletion mutant with CIP at a high concentration (10 µg/ml), which was 1,000x higher than the minimum inhibitory concentration (MIC).

Intracellular ATP concentrations were determined over six hours. In wild-type cultures, a ∼1.7-fold drop of ATP was only observed after four hours of CIP, while ATP concentrations even significantly increased in the *tisB* deletion mutant (Fig. 6A). It should be noted that the drop of ATP in CIP-treated wild-type cultures was not comparable to the drastic ATP depletion observed upon *tisB* expression from plasmid p0SD-*tisB* (Fig. 5A). However, ∼66% of wild-type cells displayed one or two IbpA-msGFP foci after six hours of CIP, indicating protein aggregation, which was not observed in the *tisB* deletion background (Fig. 6B and C). This led us to conclude (i) that TisB-dependent protein aggregation occurs in wild-type cells after prolonged DNA-damage stress, and (ii) that ATP is probably not the only determining factor for TisB-dependent protein aggregation.

**Figure 6.**
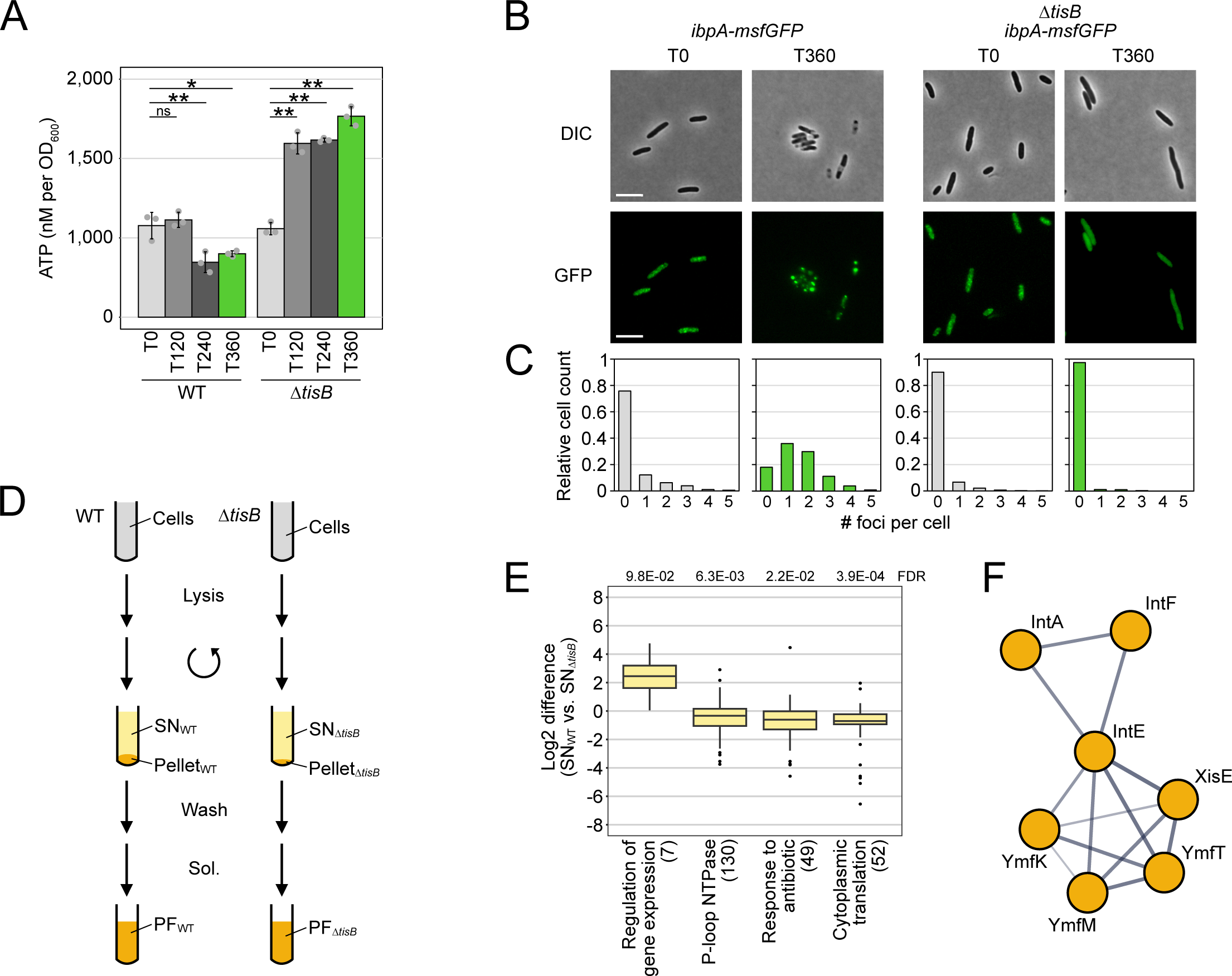
Analysis of TisB-dependent protein aggregates in wild-type cultures upon CIP treatment. **(A)** Wild type (WT) MG1655 and a *tisB* deletion mutant were treated with CIP (10 µg/ml; 1,000x MIC) during exponential phase (OD_600_ ∼0.4) for 6 hours. A luciferase-based assay was applied to measure cellular ATP levels (nM per OD_600_) before (T0) and after 120 min (T120), 240 min (T240) and 360 min (T360) of L-ara treatment. Bars represent the mean of three biological replicates, with two technical replicates each, and error bars indicate the standard deviation. Dots show individual data points. ANOVA with post-hoc Tukey HSD test was performed (* p<0.05, ** p<0.01, ns: not significant). **(B)** Strain MG1655 *ibpA-msfGFP* and Δ*tisB ibpA-msfGFP* were treated with CIP (10 µg/ml; 1,000x MIC) during exponential phase (T0; OD_600_ ∼0.4) for 6 hours (T360). Differential interference contrast (DIC) images are displayed together with corresponding fluorescence images (GFP). White bars represent a length scale of 2 µm. Representative images from three biological replicates are shown. **(C)** Quantification of msGFP foci. The experiment shown in (B) was performed in three biological replicates. All images were evaluated using a U-Net neural network analysis and in-house image processing tools to automatically count msGFP foci per cell. At least 577 cells were analyzed for each condition (*ibpA-msfGFP* T0: n=766; *ibpA-msfGFP* T360: n=577; Δ*tisB ibpA-msfGFP* T0: n=1621; Δ*tisB ibpA-msfGFP* T360: n=901). **(D)** Schematic representation of the protein aggregate purification procedure. Wild type (WT) MG1655 and a *tisB* deletion mutant (Δ*tisB*) were treated with CIP (10 µg/ml; 1,000x MIC) during exponential phase (OD_600_ ∼0.4) for 6 hours. After cell lysis and centrifugation, supernatants (SN) were collected for LC-MS analysis. The pellet fractions were washed three times and solubilized (Sol.) to receive pellet fractions (PF) for LC-MS analysis. **(E)** 1D annotation enrichment results of differentially abundant proteins in the SN_WT_ versus SN_Δ*tisB*_ (number of enriched terms in brackets; Benjamini-Hochberg FDR provided on top). **(F)** Protein-protein association network of prophage proteins in TisB-dependent protein aggregates (TdPA). The network was revealed by a multi-protein search using the STRING database (64), indicating the enrichment of the corresponding proteins within the category ‘viral process and bacteriophage tail fiber assembly’. The thickness of connecting lines indicates the confidence of the respective interaction.

### Analysis of proteomic changes and protein aggregates

We were curious about the proteins that were present in TisB-dependent protein aggregates. To this end, aggregates were purified from wild-type cultures after six hours of CIP treatment according to an established procedure (59) with minor modifications. In parallel, the *tisB* deletion mutant was analyzed as control for the identification of TisB-independent effects. After cell lysis and clearance of lysates *via* centrifugation, supernatants contained major cellular protein fractions, while pellets contained cell debris and, potentially, protein aggregates. After washing and solubilization of pellets, supernatants (SN) and pellet fractions (PF) were analyzed (Fig. 6D). The approach was initially validated by Western blot analysis using the *ibpA-msGFP* reporter background and detection of IbpA-msGFP. The Western blot corroborated our microscopic observations (Fig. 6B) and confirmed that the procedure was suitable to specifically enrich protein aggregates (Fig. S8). We then performed the experiment in wild type MG1655 and the corresponding *tisB* deletion mutant and analyzed SN and PF samples by liquid chromatography-mass spectrometry (LC-MS). We compared SN samples by label-free quantification, which allowed us to analyze the TisB-dependent response to CIP on the proteome level. Enrichment analysis revealed four functional categories that showed either increased or decreased protein abundance in the wild type as compared to the Δ*tisB* mutant (Fig. 6E). Among the category with increased protein abundance (‘regulation of gene expression’), we found several cold-shock proteins (CspA, CspC, CspD, and CspE). The remaining categories contained proteins with decreased abundance, including 130 P-loop NTPases, 49 proteins with a potential role in response to antibiotic, and 52 ribosomal subunit proteins (‘cytoplasmic translation’). We then compared the PF samples and identified 29 proteins that were significantly enriched in wild-type PF samples in comparison to Δ*tisB* (log_2_ fold-change >1 and Welch’s *t*-test with Benjamini-Hochberg FDR <0.05; Dataset S2). The sHSPs IbpA (log_2_ fold-change of 5.6) and IbpB (log_2_ fold-change of 3.2) were among the proteins with the highest enrichment factor, which confirmed successful purification of protein aggregates in wild-type PF samples. Furthermore, we identified 102 proteins that were only present in wild-type PF samples but absent from Δ*tisB* PF samples (Dataset S2). The combination of both groups (131 proteins in total) was defined as ‘TisB-dependent protein aggregates’ (TdPA). As reference for downstream analyses, the combined supernatant (cSN) of wild-type and Δ*tisB* cultures was used, which comprised 1,956 proteins in total (Fig. S8). TdPA proteins did not show a generally increased abundance in wild-type cells as compared to Δ*tisB* (Dataset S2), excluding the possibility that these proteins were solely aggregated due to their higher abundance. Furthermore, there was no intriguing difference between TdPA and cSN proteins concerning molecular weight or isoelectric point (Fig. S8). We speculated that TisB interferes with the export of outer membrane proteins (OMPs) and/ or membrane insertion of inner membrane proteins (IMPs), which would potentially cause their accumulation and aggregation in the cytoplasm (63). However, there was no clear enrichment of OMPs or IMPs in the TdPA dataset (Fig. S8). In addition, *in vitro* experiments revealed that inner membrane vesicles from CIP-treated wild- type cultures were not compromised in transport of the outer membrane protein OmpA in comparison to non-treated wild-type cultures (Fig. S8). We concluded that accumulation of membrane proteins in the cytoplasm could not account for the increased protein aggregation in CIP-treated wild-type cells. To further analyze the TdPA dataset, a STRING database search was applied (64). Protein-protein association networks and functional enrichment analyses revealed a group of seven functionally associated proteins that are encoded in the *E. coli* K-12 cryptic prophages, which applies to integrases IntA, IntF and IntE, excisionase XisE, repressor YmfK, cell division inhibitor YmfM, and transcriptional regulator YmfT (Fig. 6F). Whether these prophage proteins are related to the formation of aggregates remains unclear and needs further investigation.

### Protein aggregates determine dormancy depth of persister cells after ciprofloxacin treatment

We asked the question whether protein aggregation affects the state of persister cells upon treatment with CIP. Since the *tisB* deletion mutant does not from protein aggregates at the regular incubation temperature of 37°C, we applied heat stress at 46°C to induce aggregation (Fig. 7A). After six hours of CIP treatment at 37°C, survival was reduced by ∼20-fold in Δ*tisB* as compared to the wild type (Fig. 7B). This is in agreement with previous results showing that TisB is an important factor for persister cell survival upon CIP treatment (25, 31). At 46°C, however, survival was comparable between both strains (Fig. 7B). When applying the ScanLag method for cultures that were treated with CIP at 37°C, we observed that colonies of the Δ*tisB* mutant appeared on average 300 min earlier than wild-types colonies (Fig. 7C), indicating a reduced dormancy depth of Δ*tisB* cells, probably because aggregates were absent. When CIP was applied at 46°C, colony appearance times were comparable, suggesting that heat-induced protein aggregation has the potential to shift Δ*tisB* persisters to later appearance times by pushing them into a deeper state of dormancy.

**Figure 7.**
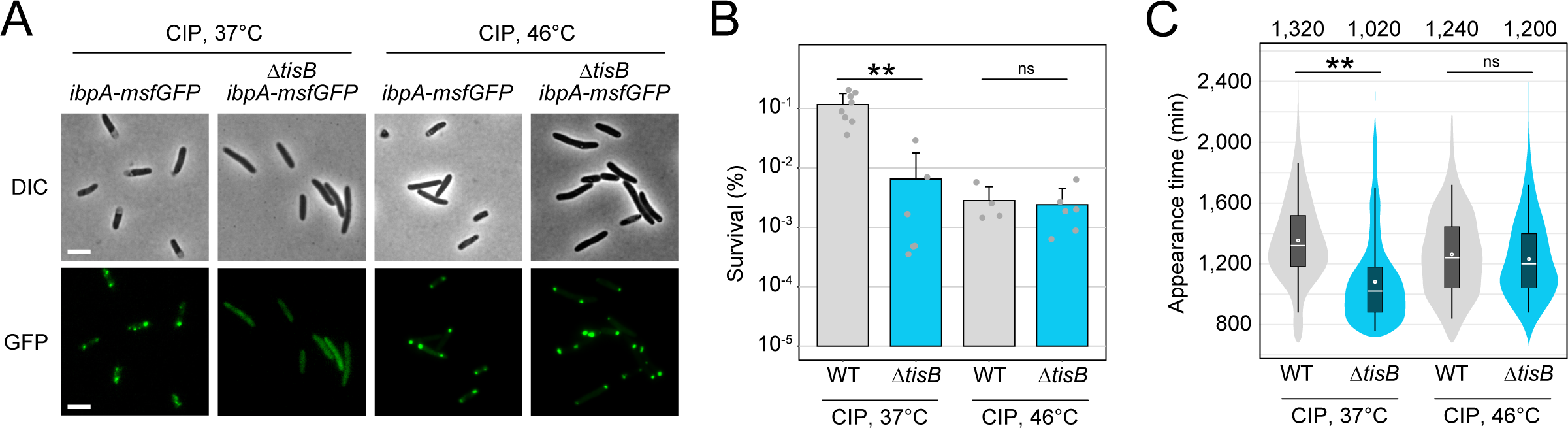
Heat-induced protein aggregates affect recovery from CIP. **(A)** Strain MG1655 *ibpA-msfGFP* and Δ*tisB ibpA-msfGFP* were treated with CIP (10 µg/ml; 1,000x MIC) during exponential phase (OD_600_ ∼0.4) for 6 hours at 37°C or 46°C. Differential interference contrast (DIC) images are displayed together with corresponding fluorescence images (GFP). White bars represent a length scale of 2 µm. **(B)** Wild type (WT) MG1655 and a *tisB* deletion were treated with ciprofloxacin (10 µg/ml; 1,000x MIC) during exponential phase (OD_600_ ∼0.4) for 6 hours at 37°C or 46°C. Pre- and post-treatment samples were used to determine relative CFU (%) by plating and counting. Bars represent the mean of at least four biological replicates and error bars indicate the standard deviation. Dots show individual data points (WT 37°C: n=8; Δ*tisB* 37°C: n=6; WT 46°C: n=4; Δ*tisB* 46°C: n=6). ANOVA with post-hoc Tukey HSD was performed (** p<0.01, ns: not significant). **(C)** Wild type (WT) MG1655 and a *tisB* deletion mutant were treated with ciprofloxacin (10 µg/ml; 1000x MIC) during exponential phase (OD_600_ ∼0.4) for 6 hours at 37°C or 46°C. ScanLag was applied to determine the colony appearance time after CIP treatment. Colony appearance times are illustrated as violin box plots. Colonies from at least three biological replicates were combined (WT 37°C: n=1431; Δ*tisB* 37°C: n=272; WT 46°C: n=476; Δ*tisB* 46°C: n=1026). The white dot indicates the mean. The respective median appearance time (white bar) is shown on top of each plot. The Δ*tisB* mutant was compared to the corresponding wild type MG1655 using a pairwise Wilcoxon rank sum test (** p<0.0001, ns: not significant).

## Discussion

Dormancy is an efficient strategy to survive harmful situations. It is, therefore, not surprising that microorganisms have evolved different mechanisms to induce dormancy. A hallmark of toxins from chromosomal TA systems is their ability to halt cell growth, induce dormancy, and eventually promote persistence, especially when toxins are expressed from plasmids (25, 30, 31, 65–68). However, strong toxin expression from plasmids does not necessarily reflect the natural situation, potentially limiting the validity of obtained effects. Here, we introduce an optimized and inducible *tisB* expression system that avoids toxic effects but retains the dormancy-promoting feature. Instead of manipulating transcription initiation (36, 48), we manipulated translation initiation by introducing an artificial SD-free 5’ UTR to the *tisB* gene on the pBAD expression plasmid. In *E. coli*, native transcripts without canonical SD sequences are not necessarily compromised in translation efficiency, suggesting that a SD sequence is not mandatory for efficient translation initiation (69). Here, we observed that the artificial SD-free 5’ UTR reduced TisB protein levels by ∼10-fold in comparison to the native *tisB* 5’ UTR. We suggest that the SD-free 5’ UTR used in this study is a valuable genetic element that enables moderate expression of toxic genes, which may be especially useful when the resulting proteins have the potential to cause cell lysis or DNA damage (70–72).

The optimized *tisB* expression system was applied to reveal the response to TisB-mediated stress in *E. coli*. Our RNA-seq data are in good agreement with an earlier transcriptome study of a *tisB* overexpression strain (44). When comparing both analyses, the most prominent up-regulated features are (i) the oxidative stress regulator gene *soxS*, (ii) the *ibpAB* operon, and (iii) CpxR-dependent genes, such as the chaperone genes *spy* and *cpxP*. It has been demonstrated that TisB provokes formation of the reactive oxygen species superoxide, leading to strong *soxS* induction (37). The ability to detoxify superoxide by the superoxide dismutases SodA and SodB is important for recovery from TisB-induced dormancy (37). Here, we observed a similar pattern: the absence of stress-related proteins (e.g., chaperones IbpB, CpxP, or Spy) delayed the recovery from TisB-induced dormancy. We conclude that the TisB-dependent stress response mainly promotes the recovery process by repairing damages and restoring cellular integrity, as we already speculated earlier (46).

Chaperones are universal to all living cells and play important roles in protein quality control and disaggregation of protein aggregates (73, 74). The sHSPs IbpA and IbpB are chaperones that initiate the disaggregation process in the cytoplasm. Further components with a pivotal role in disaggregation and ATP-dependent protein re-folding are the DnaK-DnaJ-GrpE chaperone system, the ClpB disaggregase, chaperonins GroES and GroEL, and the ATP-dependent protease HslUV. Besides *ibpAB*, both our RNA-seq approach and proteome analysis revealed TisB-dependent up-regulation of *clpB*, *groL* and *hslU*, albeit they did not match our cutoff criteria. The prevalence of chaperone genes led to the hypothesis that *tisB* expression provokes protein aggregation, and indeed, cytosolic aggregates were observed upon *tisB* expression using an established fluorescent reporter system. Besides cytosolic chaperones, our data highlight the functional importance of the periplasmic chaperones Spy and CpxP, both belonging to the CpxR regulon. While Spy is an ATP-independent chaperone that protects OMPs from folding stress (75, 76), CpxP might have a dual function by both regulating the Cpx response and acting as a chaperone (54, 77). Although not further investigated here, we suggest that *tisB* expression leads to protein folding stress in the cell envelope, thereby activating the Cpx response.

The membrane toxin TisB is well-studied with regard to its inducing condition (i.e., SOS response following DNA damage) (25, 31, 39). TisB-dependent effects can, therefore, be revealed upon treatment with the DNA-damaging antibiotic CIP (29). Antibiotics have already been associated with an increased abundance of heat shock proteins and chaperones, as e.g. observed in *Pseudomonas aeruginosa* treated with the aminoglycoside tobramycin (78), *Streptococcus pneumoniae* treated with the β-lactam penicillin (79), or *Acinetobacter baumannii* treated with different classes of antibiotics (80). In *E. coli*, deletion of heat shock proteins and chaperones resulted in a reduced survival upon treatment with levofloxacin (81), a fluoroquinolone (FQ) antibiotic that is functionally related to CIP. The authors assumed that FQ antibiotics induce the formation of cytosolic protein aggregates, which need to be disassembled by heat shock proteins and chaperones (81). Intriguingly, we were able to demonstrate that protein aggregation occurs upon treatment with CIP, and that this process depends on TisB, suggesting that TisB is the foremost factor for protein aggregation in response to FQ-induced DNA damage. The question remains how a membrane toxin provokes aggregation. Hypothetically, TisB accumulates in the cytoplasm and initiates a nucleation process that leads to aggregate formation (82). However, cellular fractionation experiments combined with Western blot analysis indicate that TisB does not accumulate in the cytoplasm but rather completely localizes to the membrane (our unpublished results). Furthermore, we show here that production of the attenuated toxin TisB-K12L does not cause aggregation. We conclude that aggregation is a downstream effect of TisB and its function as a pore-forming toxin. The primary action of TisB is breakdown of the proton motive force, which is similar to the action of protonophores and likely leads to disturbance of pH homeostasis and acidification of the cytoplasm (83, 84). Potentially, the drop in intracellular pH initiates the aggregation process (85), but this needs further investigation. Interestingly, pH-driven protein aggregation and acidification have been associated with dormancy and increased stress tolerance in both yeast and bacteria (86, 87), and may also contribute to TisB-induced dormancy and persistence.

Our proteome study revealed that many P-loop NTPases showed a decreased abundance in CIP-treated wild-type cells as compared to the *tisB* deletion strain. These enzymes utilize energy from ATP (or GTP) hydrolysis (88), and their decreased abundance might be a reaction to the reduced energy levels observed in TisB-producing cells. In addition, many ribosomal subunit proteins had a decreased abundance. Collectively, these findings indicate that TisB-producing cells reduce energy-consuming core processes, such as replication and translation, which might avoid further DNA damage by CIP and support dormancy as such. Analysis of TisB-dependent protein aggregates revealed the enrichment of proteins from cryptic prophages, and it remains an open question whether this is coincidental or has a biological meaning. *E. coli* K-12 harbors nine cryptic prophages, and – albeit their functions remain largely unknown – it has been observed that they contribute to survival under antibiotic stress, including the DNA-damaging quinolone nalidixic acid (89). Potentially, the prophage proteins contribute to the aggregation process, which in turn affects antibiotic tolerance and persistence.

Finally, we propose that the primary function of the membrane toxin TisB is the establishment of dormant cells through energy depletion, but that secondary effects determine the dormancy depth. In this study, we revealed protein aggregation as one of the most important factors for dormancy depth in TisB-expressing cells. This is in line with previous observations that protein aggregation is correlated with dormancy depth (58) and may even drive bacteria into a state of deep dormancy, such as the viable but nonculturable (VBNC) state (59). It remains an exciting task for future studies to reveal the molecular mechanisms behind the TisB-induced protein aggregation process.

## Materials and Methods

### Growth conditions

*E. coli* strains were grown in lysogeny broth (LB) medium at 37°C and 180 rpm. If temperature-sensitive plasmids were present, strains were grown at 30°C and 180 rpm. Pre-cultures were cultivated in the presence of antibiotics if applicable (50 µg/ml kanamycin, 15 µg/ml chloramphenicol, 200 µg/ml ampicillin and 6 µg/ml tetracycline). Pre-cultures were diluted 100-fold into fresh LB medium and grown until exponential phase was reached. Growth curves were recorded in 30-min time intervals with a cell density meter model 40 (Fisher Scientific).

### Construction of strains and plasmids

*E. coli* strains used in this study are derivatives of K-12 wild type MG1655 and are listed in Table S1. Chromosomal deletion mutants were constructed using the λ red methodology (90). A selection marker (*cat* or *kan* gene) was amplified via PCR using primers with specific 40-bp overhangs, matching the desired deletion locus. An *E. coli* MG1655 strain, that provides the heat-inducible λ red genes on plasmid pSIM5-tet (91), was grown at 30°C in the presence of tetracycline (3 µg/ml) until an OD_600_ of ∼0.4 was reached. After a 15-min heat shock at 42°C, electrocompetent cells were prepared and PCR products were transformed *via* electroporation. Clones were selected on LB agar plates containing the appropriate antibiotic (12.5 µg/ml chloramphenicol or 50 µg/µl kanamycin), and gene deletions were subsequently verified by colony PCR. After two incubations at 42°C, loss of the heat-sensitive plasmid pSIM5-tet was verified by tetracycline sensitivity. If necessary, deletion constructs were transduced into new strain backgrounds using P1 phages according to standard protocols.

Expression plasmid p0SD-*tisB* was generated by AQUA cloning (92). The *tisB* insert was amplified by PCR using primer pair AQ-0ATG-2-f/ NES-rev and plasmid p+42-*tisB* as template. Primer AQ-0ATG-2-f provides both a 20-bp artificial 5’ UTR (lacking a Shine-Dalgarno sequence) and a 20-bp overhang for AQUA cloning. The pBAD backbone was amplified by PCR using primer pair AQ-topo-f/ AQ-topo-rev to generate matching overhangs for the *tisB* insert. Purified amplification products were mixed in a final volume of 10 µl, applying a molecular ratio of 7:1 (insert to backbone; 100 ng backbone). Mixtures were incubated at 25°C for one hour. Afterwards, mixtures were transformed into chemically competent MG1655 cells and clones were selected on LB agar containing ampicillin (200 µg/ml). In a similar way, plasmid p0SD-*3xFLAG-tisB* was generated with primer pair AQ-0ATG-3x-f/ NES-rev using plasmid p+42-*3xFLAG-tisB* as template for amplification of the *3xFLAG-tisB* insert. Plasmid p0SD-*tisB- K12L* was generated by site-directed mutagenesis PCR using primer pair K12L-for/ K12L-rev and plasmid p0SD-*tisB* as template. After PCR, parental plasmids were digested with DpnI (Thermo Fisher Scientific) and the linear PCR product was transformed into chemically competent MG1655 cells. Clones were selected on LB agar containing ampicillin (200 µg/ml). All plasmids were verified by Sanger sequencing (Microsynth SeqLab, Göttingen, Germany) and are listed in Table S2. Primers used for cloning procedures are listed in Table S3.

### Determination of relative colony counts and persister levels

Exponential-phase cultures (OD_600_ ∼0.4) were treated with L-ara (0.2%) for one hour or with CIP (10 µg/µl; 1,000x MIC) for six hours at 37°C and 180 rpm. Pre- and post-treatment samples were serially diluted and plated on LB agar plates. In case of L-ara treatment, cells were diluted with NaCl (0.9%). In case of CIP treatment, cells were washed two times and diluted with 20 mM MgSO_4_. Colonies were counted after ∿20 hours (pre-treatment) or ∿40 hours (post-treatment). Colony counts were used to determine colony forming units (CFU) per milliliter. The ratio between treated and untreated samples represents either the relative CFU count (L-ara) or persister level (CIP). P-values were calculated using an ANOVA with a post-hoc Tukey HSD test in R statistical language (https://www.r-project.org/).

### Analysis of colony growth

Colony growth was analyzed using the ScanLag method (52). Agar plates from L-ara or CIP treatments (see *Determination of relative CFU counts and persister levels*) were covered with black felt, placed on scanners and incubated at 37°C. Epson Perfection V39 scanners were used to record time series of images controlled by the *ScanningManager* application. Images (TIFF files) were taken every 20 min for a total period of 40 hours. Image processing was performed using MatLab (MathWorks) with functions *PreparePictures, setMaskApp, TimeLapse* and *ScanLagApp* (51). After image processing, the appearance and growth times were extracted. The appearance time is defined by a colony size of 10 pixels, whereas the growth time is defined as the time that is needed to cause a colony size increase from 80 to 160 pixels. Growth data was used to create violin box plots with Power BI Desktop (Microsoft). P-values were calculated using a pairwise Wilcoxon rank sum test in R statistical language (https://www.r-project.org/).

### Membrane depolarization measurements

Exponential-phase cultures (OD_600_ ∼0.4) were treated with 0,2% L-ara for one hour at 37°C and 180 rpm. Samples were withdrawn before and after addition of L-ara and adjusted to an OD_600_ of 0.5. DiBAC_4_(3) (Sigma-Aldrich) was added at a final concentration of 1 µg/ml, followed by incubation for 20 min in the dark at room temperature. Cells were washed two times with 200 µl 1x PBS at 13,000 rpm for 3 min at room temperature and finally resuspended in 250 µl 1x PBS. DiBAC_4_(3) fluorescence (excitation: 490 nm, emission: 520 nm) was measured in technical duplicates (100 µl each) in an Infinite M Nano^+^ microplate reader (Tecan). P-values were calculated using an ANOVA with a post-hoc Tukey HSD test in R statistical language (https://www.r-project.org/).

### ATP measurements

Cultures were grown to exponential phase (OD_600_ ∼0.4) and treated with L-ara (0.2%) for one hour or with CIP (10 µg/µl; 1,000x MIC) for up to six hours. Samples (1 ml) were withdrawn before and after treatment. Cell pellets were collected by centrifugation (13,000 rpm, 3 min) and supernatants were discarded. Cells were washed with 1 ml NaCl (0.9%) and resuspended in 1 ml LB medium. 100 µl of samples were mixed with 100 µl BacTiter-Glo^TM^ reagent (Promega) and incubated for 5 min in the dark. The luminescence was measured using an Infinite M Nano^+^ microplate reader (Tecan). Values were transformed to nM, using the slope formula of an ATP calibration curve, and normalized to the OD_600_. P-values were calculated using an ANOVA with a post-hoc Tukey HSD test in R statistical language (https://www.r-project.org/).

### Fluorescence microscopy

Cultures were grown to exponential phase (OD_600_ ∼0.4) and treated with L-ara (0.2%) for one hour or with CIP (10 µg/µl; 1,000x MIC) for six hours at 37°C or 46°C. Samples before and after treatment were transferred onto agarose pads (1% agarose in 1x PBS) on top of a microscopy slide with a cover slip on top of the cells. Images were recorded with a Leica DMI 6000 B inverse microscope (Leica Camera AG) using an HCX PL APO 100x/1.4 differential interference contrast (DIC) objective, a pco.edge sCMOS camera (PCO AG), and software VisiView version 4.3.0 (Visitron Systems GmbH). For fluorescence images (GFP), a custom filter set (T495lpxr, EX470/40m; EM525/50; Chroma Technology) was used. The exposure time was set to 50 ms with a binning of 2 and an offset of 0.0. Images were saved as TIFF and further processed with the open-source software ImageJ version 1.53k.

### Automated foci analysis

For U-Net training and segmentation, DIC images of *E. coli* cells were used. The software used was the U-Net plugin for ImageJ, available from the website of the Computer Vision Group at the University of Freiburg (61). For training, 906 cells in eight images were annotated. To enhance segmentation quality and facilitate separation of cell aggregates into individual cells, one label was used for the circumference of the cells and one for their inside. A training with 2000 iterations and learning rate of 1E-4 yielded segmentations that were very close to the training annotation. With post-processing using a custom Wolfram Mathematica script, the segmentations were further refined and crooked, very small, very large features or cells at the image border were excluded. Visual inspection of all segmentations confirmed that the vast majority of cells were properly identified. The extracted cell shapes were used as masks for the GFP image channel. Spatial filtering, peak finding and thresholding yielded the foci.

### Preparation of RNA-sequencing samples

Exponential-phase cultures (OD_600_ ∼0.4) of strain MG1655 p0SD-*tisB* were treated with 0,2% L-ara to induce *tisB* expression for 30 min. Samples from biological triplicates were withdrawn before (samples “Exp”) and after L-ara treatment (samples “T30”) and immediately inactivated by adding 200 µl stop solution (95% ethanol, 5% phenol) to 1 ml cell culture on ice. Total RNA was isolated according to the hot acid-phenol method as described (38). DNA was removed using the TURBO DNA-*free*^TM^ kit (Invitrogen) according to the ‘rigorous treatment’ instructions. The final clean-up was performed using phenol/chloroform/isoamyl alcohol (25:24:1) mixed with the sample in a 1:1 ratio, followed by chloroform treatment and precipitation as before. RNA quality was assessed on an 8% polyacrylamide gel containing 1x TBE and 7 M urea. Aliquots of approximately 3.5 µg of total RNA were prepared and stored at -80°C until further analysis.

### RNA-sequencing and data analysis

RNA-sequencing was performed by vertis Biotechnologie AG. For cDNA synthesis, all RNA samples were first fragmented using ultrasound (4 pulses of 30 seconds, each at 4°C). Then, an oligonucleotide adapter was ligated to the 3’ end of the RNA molecules. First-strand cDNA synthesis was performed using M-MLV reverse transcriptase and the 3’ adapter as primer. The first-strand cDNA was purified and the 5’ Illumina TruSeq sequencing adapter was ligated to the 3’ end of the antisense cDNA. The resulting cDNA was PCR-amplified to about 10-20 ng/µl using a high-fidelity DNA polymerase for 12 cycles. The TruSeq barcode sequences, which are part of the 5’ and 3’ TruSeq sequencing adapters, were used. The cDNA was purified using the Agencourt AMPure XP kit (Beckman Coulter Genomics) and analyzed by capillary electrophoresis. For Illumina NextSeq sequencing, the samples were pooled in approximately equimolar amounts. The cDNA pool in the size range of 200-550 bp was eluted from a preparative agarose gel. An aliquot of the size fractionated pool was analyzed by capillary electrophoresis. The cDNA pool was single-read sequenced on an Illumina NextSeq 500 system using 75-bp read length.

Quality and adapter trimming was performed with Trim Galore (Version 0.6.5) (https://github.com/FelixKrueger/TrimGalore) with Cutadapt Version 2.7 (http://dx.doi.org/10.14806/ej.17.1.200) using the parameters ‘--quality 20 --length 20’ and default adapter detection and trimming. MultiQC (Version 1.8) (93) and FastQC (Version 0.11.8) (http://www.bioinformatics.babraham.ac.uk/projects/fastqc/) were used for quality control. The preprocessed reads were aligned with Bowtie2 (Version 2.3.5) (94) using the ‘--mm’ and ‘--very-sensitive’ settings and GCF_000005845.2 (NCBI; downloaded 25.11.2019) as a reference genome. For postprocessing of the alignments, gene counting and data analysis, Samtools (Version 1.9) (95), featureCounts (Version 1.6.4) (96) and DESeq2 (Version 1.26) (97) were applied, respectively. All bioinformatic calculations were performed using Curare (Version 0.1.1) (https://github.com/pblumenkamp/Curare) and R statistical language (https://www.r-project.org/). Processed RNA-seq data are available as Dataset S1 and have been deposited together with raw data files on the NCBI Gene Expression Omnibus (GEO) under the accession number GSE255764.

### Northern blot analysis

Cultures were grown to exponential phase (OD_600_ ∼0.4) and treated with L-ara (0.2%) for 30 min. Total RNA for Northern blot analysis was isolated using the hot acid-phenol method as described (38). Northern blot analysis was performed with 5-10 µg of total RNA. The RNA was separated using 10% polyacrylamide gels containing 1x TBE and 7 M urea at 300 V for approximately three hours. The RNA was transferred to a Roti®Nylon plus membrane (Roth) by semi-dry electroblotting at 250 mA for three hours. After UV-crosslinking, the membrane was pre-hybridized using Church buffer [0.5 M phosphate buffer (pH 7.2), 1% (w/v) bovine serum albumin, 1 mM EDTA, 7% (w/v) SDS] at 42°C for one hour. Hybridization with probes for detection was performed overnight. Specific probes were generated by end-labeling of oligodeoxyribonucleotides (Table S3) using T4 Polynucleotide Kinase (New England Biolabs) and [γ-^32^P]-ATP (Hartmann Analytic). Membranes were washed (5x SSC, 0.01% SDS) and exposed to phosphorimaging screens (Bio-Rad). Screens were analyzed using a Molecular Imager FX and the Quantity One 1-D Analysis Software (Bio-Rad).

### Quantitative reverse transcription PCR

Cultures were grown to exponential phase (OD_600_ ∼0.4) and treated with L-ara (0.2%) 30 min. Total RNA for quantitative reverse transcription PCR (qRT-PCR) was isolated using the NucleoSpin® RNA Kit (Macherey-Nagel), including DNA digestion. RNA concentrations were measured using a spectrophotometer (NanoDrop 1000) and subsequently adjusted to a concentration of 5 ng/µl. For reverse transcription and amplification of gene-specific fragments, 10-µl reactions were prepared with the Brilliant III Ultra-Fast SYBR Green qRT-PCR Master Mix (Agilent) in technical duplicates for each sample. Reactions contained 1 ng/µl of total RNA and 0.5 µM of each primer (Table S3). Reverse transcription and amplification were performed on a CFX Connect Real-Time System (Bio-Rad). Reverse transcription was carried out at 50°C for 10 min followed by 95°C for 3 min. For amplification, 45 cycles were applied at 95°C for 5 seconds, 56°C for 10 seconds and 72°C for 10 seconds (*tisB* gene), or at 95°C for 5 seconds and 60°C for 10 seconds (all remaining genes). Amplification curves were recorded with the CFX Maestro software (Bio-Rad). Cq values were used to calculate fold-changes according to Pfaffl (98). The *hcaT* gene was used as reference for normalization (38).

### Western blot analysis

For detection of 3xFLAG-TisB, strains were grown to exponential phase. Samples were withdrawn in a defined volume (equivalent to an OD_600_ of 10) and centrifuged at 10,000 rpm and 4°C for 10 min. Cell pellets were resuspended in 50 µl SDS sample buffer (12% SDS, 6% β-mercaptoethanol, 30% glycerol, 0.05% Coomassie blue, 150 mM Tris/HCl, pH 7.0). For protein separation, a Tricine-SDS-PAGE was applied with 16% polyacrylamide (99). Samples were incubated at 95°C for 10 min before loading onto the gel. An initial voltage of 60 V was applied until samples entered the separation gel. Afterwards, electrophoresis was carried out at 100 V for about three hours. Proteins were transferred onto a PVDF membrane by semi-dry electroblotting over night at 0.4 mA/cm². Membranes were stained with Ponceau S and documented before blocking with 5% milk powder in 1x PBST (PBS + 0.1% Tween20) for one hour. For detection of 3xFLAG-TisB, membranes were incubated with an HRP-conjugated monoclonal IgG α-FLAG antibody (Sigma-Aldrich) in 3% BSA in PBST at room temperature for 90 min. Using the Lumi-Light Western Blotting Substrate (Roche), 3xFLAG-TisB was visualized and documented in a chemiluminescence imager (PeqLab) with the FusionCapt Advance software (Vilber Lourmat).

### Purification of protein aggregates

Protein aggregates were purified according to a published protocol (59) with minor modifications. Strains were grown to exponential phase (OD_600_ ∼0.4) and treated with CIP (10 µg/ml; 1,000x MIC) for six hours. Cells (38 ml culture volume) were harvested and centrifuged at 4,000 x *g* and 4°C for 30 min. Cells were resuspended in 10 ml washing buffer I (300 mM NaCl, 5 mM β-mercaptoethanol, 1 mM EDTA, 50 mM HEPES, pH 7.5) and centrifuged as before. The cell pellet was dissolved in 10 ml lysis buffer (washing buffer I containing 1 µg/ml leupeptin and 0.1 mg/ml AEBSF [4-(2- aminoethyl)benzenesulfonyl fluoride hydrochloride]). Cells were lysed in three cycles with a cell homogenizer at 1,380 to 1,725 bar, followed by centrifugation at 11,000 x *g* and 4°C for 30 min to clear the lysates. Supernatants (SN) were stored at -80°C until LC-MS analysis. Pellets were resuspended in 2 ml washing buffer II (washing buffer I containing 0.8% Triton X-100 and 0.1% sodium deoxycholate) and centrifuged as before. The washing step was repeated two more times. After the final washing step, pellets were resuspended in 1 ml solubilization buffer (1% SDS, 1x SigmaFast Protease Inhibitor [Sigma-Aldrich], 50 mM HEPES, pH 8.0). Pellet fractions (PF) were stored at -80°C until LC-MS analysis.

### LC-MS based proteome analysis

Samples generated via the purification of protein aggregates, i.e. lysate supernatants and the protein aggregate pellets, were processed following the SP3 protocol (100). For the lysate supernatants and protein aggregate samples, 50 μg of each was analyzed. Briefly, all samples (in triplicate) were resuspended in 75 μl of 100 mM ammonium bicarbonate (ABC) buffer (pH 7.4). Samples were reduced in the presence of tris(2-carboxyethyl)phosphine (5 mM) (1 hour, 56°C), before alkylation was performed with chloroacetamide (50 mM) (room temperature (RT) in the dark, 30 min). Beads (SpeedBeads Magnetic Carboxylate) were washed twice with Milli-Q water, and 100 μg of beads in 250 mM ABC buffer were added to each sample (final volume of 100 μl per sample). Precipitation of the proteins onto the beads was initiated *via* the addition of 100 μl of ethanol, the samples were gently shaken (5 min, 800 rpm) before a further 300 μl of ethanol was added and the samples were gently shaken (800 rpm) for an additional 20 min (final concentration of ca. 80% ethanol). The bead-associated precipitated proteins were pelleted by centrifugation (21,100 x *g*, 5 min, RT) with magnet-assisted isolation to assist aspiration of the solution. The beads were then washed twice with 80% ethanol, with centrifugation (21,100 x *g*, 5 min, RT) and magnet-assisted aspiration to remove all liquid. The samples were briefly sonicated in a sonication bath between washes to aid in the re- solubilization of the protein-associated beads. Following the final wash, the beads were suspended in 100 μl of 100 mM ABC buffer containing trypsin (0.4 μg in total per sample, enzyme to protein ratio of 1:125), the samples were briefly sonicated to ensure no aggregation of the beads, then incubated overnight (37°C, shaking at 1,300 rpm). Following overnight digestion, the samples were centrifuged (21,100 x *g,* 5 min), before magnet-assisted collection of the peptide-containing supernatant was performed. The peptides were cleaned up *via* solid phase extraction (SPE) using Pierce C18 Tips 100 μl (as per manufactures protocol). Following clean-up, the supernatants were dried down *via* vacuum centrifugation and stored at -20°C. On the day of MS analysis, peptides were resuspended in 20 μl of HPLC loading buffer (3% acetonitrile and 0.1% trifluoroacetic acid).

Chromatographic separation was performed on a Dionex U3000 Nano-HPLC system equipped with an Acclaim^TM^ PepMap^TM^ 100 C18 column (2 μm particle size, 75 μm x 500 mm) coupled online to a mass spectrometer. The eluents used were: eluent A (0.05 % formic acid) and eluent B (80% acetonitrile and 0.04% formic acid). The separation was performed over a programmed 120-min run. Initial chromatographic conditions were 4% eluent B for 4 min followed by linear gradients from 4 to 50% eluent B over 90 min, then 50 to 95 % over 8 min, and 8 min at 95% eluent B. Following this, an inter-run equilibration of the column was performed (20 min at 4% eluent B). A 300 nl/min flow rate and 1 μl of sample were injected per run. Two wash runs (loading buffer injections) were performed between each sample. Data acquisition following separation was performed on a Q Exactive^TM^ Plus mass spectrometer (Thermo Fisher Scientific). A full scan MS acquisition was performed (350-1000 m/z, resolution 70,000) with the subsequent data-dependent MS/MS acquisition for the top 15 most intense ions via HCD activation at NCE 26 (resolution 17,500); an isolation window of 3 m/z was employed with apex trigger (3-15 s) and dynamic exclusion (30 s duration) enabled.

Bottom-up proteomic data analysis was performed using Proteome Discoverer (Ver. 3.0.1.27) (Thermo Fisher Scientific), and the Chimerys search algorithm. Additionally, the Minora node was included to enable label-free quantification. Raw data files were searched against a protein FASTA database containing the complete UniProt *E. coli* (K-12 substrain MG1655) protein FASTA (accessed from UniProt 2023.04.11) plus the list of common laboratory contaminants (cRAP47). The searches were conducted with trypsin specificity, allowing a maximum of two missed cleavages. Strict parsimony criteria were applied with high stringency at both the protein and peptide levels (protein level false discovery rate (FDR) <1%), and at least one high unique confidence peptide (PSM level FDR <1%). Statistical assessment of the data was performed using the Perseus software package (Ver. 2.0.10.0). The Welch’s *t*-test was performed with a minimum of two valid quantification values required for each protein in both groups, and Benjamini-Hochberg FDR calculation was performed at both medium (FDR <5%) and high (FDR <1%) cut-off levels. In addition, an abundance fold-change of greater than 2 (i.e., log_2_ difference <-1 or >1) was required. Further assessment of potentially enriched protein categories was performed *via* 1D annotation enrichment (Benjamini-Hochberg FDR <0.1) for the supernatant (SN) samples. The mass spectrometry proteomics data have been deposited to the ProteomeXchange Consortium (101) *via* the PRIDE partner repository with the dataset identifier PXD049478.

### Bioinformatics data analysis

For bioinformatics data analysis of protein aggregates, two different datasets were defined. All proteins, that were identified in at least two biological replicates of either wild-type or Δ*tisB* supernatant samples, were used as reference and referred to as combined supernatant (cSN). All proteins, that were exclusively present or enriched in wild-type pellet fractions in comparison to Δ*tisB* pellet fractions, were defined as TisB-dependent protein aggregates (TdPA). For prediction of protein localization, LocTree3 was used (102). The file 83333_Escherichia_coli.bact.lc3 was retrieved from Bacteria.zip and used to assign the localization to each identified protein. For protein-protein association networks and functional enrichment analyses, a multi-protein search in the STRING database was performed (64).

## Acknowledgments

We are grateful to Niklas Philipp for help with data analysis. We would like to thank Nasrine Bekhedda and Dana Sensen for experimental support. Abram Aertsen is acknowledged for providing the *ibpA- msGFP* reporter strain. We further thank Liliya Chernova for discussion on protein aggregation, and Kai Thormann for support with microscopy. Work in the group of B.A.B. was supported by the German Research Council (DFG) in the framework of the SPP2002 (BE 5210/3-1 and BE 5210/3-2) and by Fonds der Chemischen Industrie (material cost allowance). A.T. and L.C. were supported by DFG in the framework of the SPP2002, project Z2 (TH872/10-2). H.-G.K. was supported by the DFG in the framework of the SPP2002 (KO2184/9.1 and KO2184/9.2) and by the SFB1381 (Project ID 403222702).

